# Lesion-induced impairment of therapeutic capacities of olfactory ensheathing cells in an autologous transplantation model for treatment of spinal cord injury

**DOI:** 10.1101/2024.04.19.590121

**Authors:** Quentin Delarue, Matthis Brodier, Pauline Neveu, Laurine Moncomble, Alizée Hugede, Axelle Blondin, Amandine Robac, Clémence Raimond, Pamela Lecras, Gaëtan Riou, Nicolas Guérout

**Author notes:** **Corresponding Authors:** Dr. Quentin Delarue, Pr. Nicolas Guérout.

## Abstract

Spinal cord injury (SCI) is a serious pathology of the central nervous system that results in loss of motor, sensory and autonomic functions below the level of the lesion and for which, unfortunately, there is currently no cure. In addition to the loss of function, SCI induces a systemic inflammation that is not confined to the spinal cord and whose effects are increasingly well characterized. In particular, SCI causes cerebral inflammation, which is responsible for the impairment of hippocampal and bulbar neurogenesis. Many therapies have been tested as potential treatments for SCI. In animal models, cell therapies have shown interesting effects such as spinal scar reduction, anti-inflammatory properties, axonal regrowth or neuronal survival, allowing better functional recovery. However, in human studies, their therapeutic capacities are less significant. Beyond obvious differences in pathophysiology and cell culture procedures, a key paradigm of cell transplantation differs between humans and animals. In animal models, transplanted cells are systematically taken from healthy individuals, whereas in humans the immune incompatibility leads to the realization of autologous transplantation. Therefore, we were interested in the lesion effects on the neuro-repairing potential of olfactory ensheathing cells (OECs) harvested from olfactory bulbs.

Using functional sensory-motor studies, histological and gene expression analyses, we were able to demonstrate for the first time that the lesion negatively affects the therapeutic properties of cells used to treat SCI. These innovative results shed new light on the future use of cell transplantation in autologous transplantation after SCI.

## INTRODUCTION

Spinal cord injury (SCI) is a rare traumatic pathology of the spinal cord. It is caused by road or sports-related accidents, falls, or violence [1]. It is more common in young people and the over-60s, who are generally in good health [2]. Unfortunately, no curative treatment can be offered to people with SCI, so these patients remain disabled for life. Depending on the severity of the lesion, SCI can result in complete or incomplete motor, sensory and/or autonomic deficits below the site of injury.

From a cellular and molecular point of view, the primary traumatic lesion is followed by a secondary lesion linked to the tissue reactivity. In particular, the initial damage induces blood vessel rupture, neuronal and oligodendrocyte cell death, which leads to neuronal dysfunction and immune cells infiltration such as neutrophils and macrophages, all of which increase the reactivity of resident immune cells, microglial cells, leading to the emergence of an inflammatory response [3,4]. The reactivity of these different immune cells induces a response from astrocytes, ependymal cells and pericytes, forming a spinal scar that prevents the lesion from spreading and secondary damages caused by inflammation [5–7]. The series of events following SCI has been well described and characterized in the spinal cord. However, it has been reported that SCI is responsible for systemic inflammation [8]. The inflammatory state is also found in the brain, particularly in the hippocampus and olfactory bulb (OB), leading to a decrease in neurogenesis and cognitive decline in mice and rats [9,10].

Many treatments have been investigated in experimental or clinical trials to provide a curative therapy for these patients. These treatments are based on different paradigms, including the modulation of the lesion environment by enzymes to increase axon regrowth or, more recently, methods using human-machine interfaces [11–13]. These therapeutic approaches include cell transplantation which is one of the most widely studied. Different cell types have been studied after SCI: hematopoietic or mesenchymal stem cells [14,15] or differentiated cells such as Schwann cells [16] or olfactory ensheathing cells (OECs) [17]. Although cell types differ in origin and characteristics, identical regenerative mechanisms have been described in several studies. These cells have a therapeutic effect on SCI by modulating the spinal scar and reducing secondary damages [18].

Among cells studied, OECs are a very interesting therapeutic candidate. Their transplantation after SCI has shown anti-inflammatory and regenerative properties, allowing functional recovery in animal models. Indeed, these cells have been described to induce axonal survival and regrowth, but also to modulate the inflammatory response that occurs after SCI by acting on the polarization of microglia and macrophages, triggering a switch from a pro-inflammatory to an anti-inflammatory phenotype [19,20]. The properties of OECs and their ability to induce tissue repair and functional recovery after transplantation into SCI, without integrating into the spinal parenchyma, have led to their use in clinical trials. However, studies of bulbar OEC (bOEC) transplantation in humans have reported inconsistent results [21]. Indeed, some studies report an improvement of the ASIA score in treated patients while others show no effect after treatment.

Differences in culture, transplantation methods and physiopathology between humans and rodents may explain these changes in results. However, another important difference between animal and human studies is the transplantation paradigm. In humans, transplantation is autologous, meaning that cells come from the traumatized organism, whereas in animals, healthy animals are used for cell culture. The fact that many studies have shown that SCI can induce systemic inflammation [10,22,23]; we are wondering if the lesion itself could affect neuro-regenerative properties of the transplanted cells. Therefore, in our study, we investigated the effects of SCI on bOECs properties in the context of transplantation for SCI treatment by mimicking an autologous transplantation model in mice.

## MATERIALS AND METHODS

### Animal care and use statement

The animal protocol was designed to minimize pain or discomfort to the animals. All experimental procedures were in accordance with the European Community guidelines on the care and use of animals (86/609/CEE; Official Journal of the European Communities no. L358; 18 December 1986), the French Decree no. 97/748 of 19 October 1987 (Journal Officiel de la République Française; 20 October 1987) and recommendations of the Cenomexa Ethics Committee (#26905).

### Animals

Mice of either sex were group housed (two–five mice/cage, genders separated) in secure conventional rodent facilities on a 12-h light/dark cycle with constant access to food and water. A total of 270 mice were included in this study. We used three mouse lines in this study: 254 wild type (WT) C57BL/6 mice, 12 Rosa26-tdTomato mice and 4 (CAG-luc-GFP) L2G85Chco+/+ (FVB-Tg(CAG-luc,-GFP) L2G85Chco/J,) mice called Luciferase (LUX) thereafter. Then, LUX mice have been crossed with C57BL/6 mice three times consecutively to avoid any problem related to the strains. LUX mice were obtained from the Jackson Laboratory (https://www.jax.org/strain/008450, RRID:IMSR_JAX:008450).

All experiments were performed on adult mice aged 8-12 weeks (average weight 20 g for females and 25 g for males). All experimental groups were created with a sex ratio of 50:50. Our study consisted of four main experimental groups (Fig. 1):

**Figure 1:**
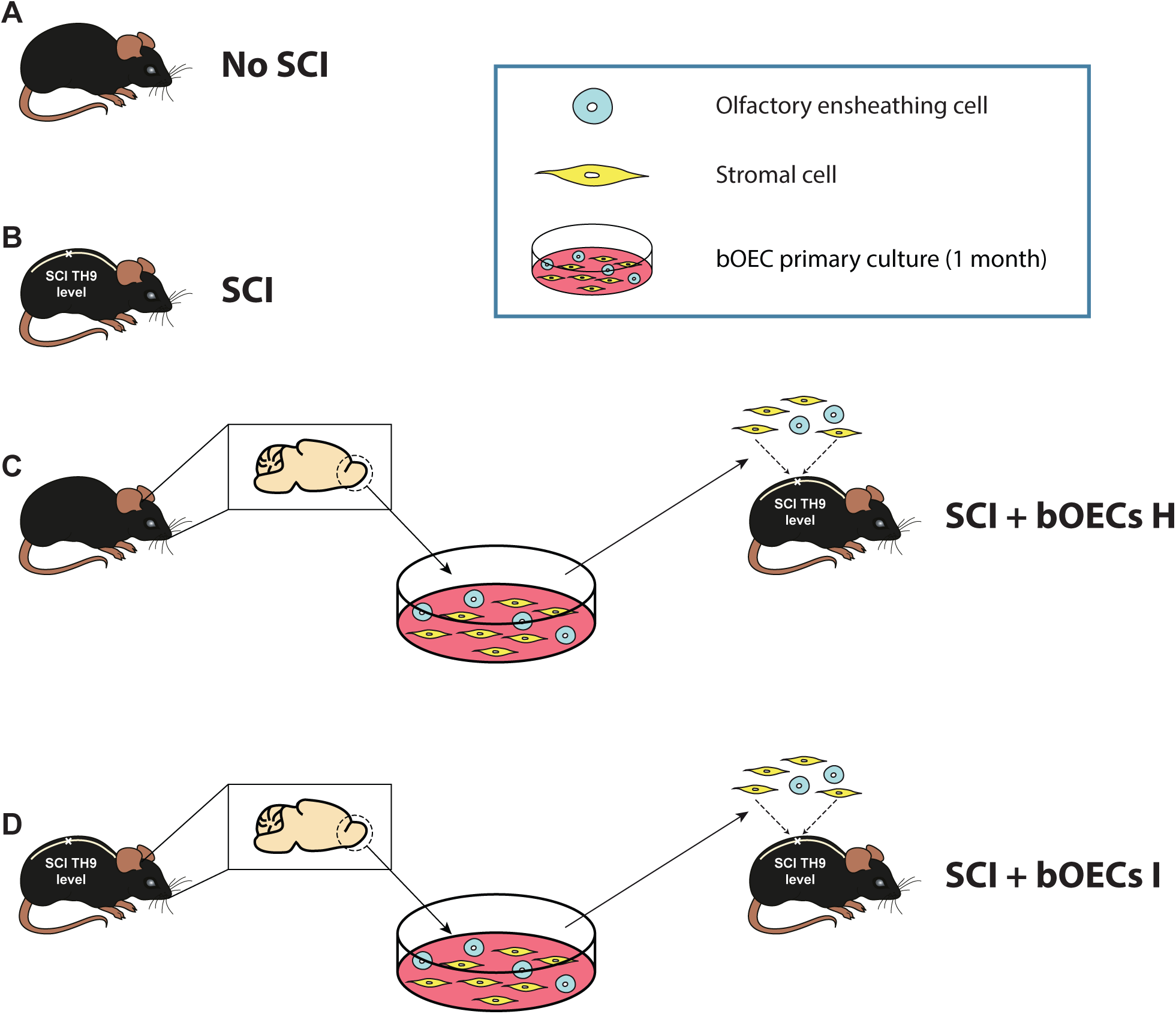
Schematic representation of the four main experimental groups used in our study. **A**. No SCI group, uninjured animals used to define the baselines for functional and histological experiments. **B**. SCI group, injured animals without treatment. **C**. SCI bOECs H, these animals underwent sci and primary bOECs were transplanted immediately. OB from uninjured animals were used to prepare bOEC cultures. **D**. SCI bOECs I, these animals underwent sci and primary bOECs were transplanted immediately. OB from injured animals were used to prepare bOEC cultures. This SCI was performed on a different set of mice 7 days prior to the primary OB cultures.

Non-injured control group: animals without SCI used to define baselines for histological and functional experiments.

SCI group: animal received SCI without transplantation.

SCI + bOECs H group: animal received SCI and primary bOECs transplantation was performed immediately after the lesion. OB used to prepare primary bOECs culture was obtained from uninjured animals.

SCI + bOECs I group: animal received SCI and primary bOECs transplantation was performed immediately after the lesion. OB used to prepare primary bOECs culture was obtained from previously SCI injured animals.

### Primary OB culture and cell transplant preparation

Briefly, mice were anaesthetized with 2% isoflurane (Iso-Vet, Osalia, Paris, France), then euthanized by decapitation, and OBs were extracted from the brain and placed in cold phosphate buffered saline (PBS). After incubation in trypsin 0.25% (2.5% Trypsin 10X, 15090-046, Thermofisher), the tissue was mechanically dissociated. After dissociation, cells were cultured in T25 flasks with 5 ml DF-10S medium (DMEM:F12 + Glutamax, 0.5% penicillin/streptomycin and 10% heat-inactivated fetal bovine serum). One month after plating, cultures were trypsinized and cells were counted to prepare a cell suspension of 25,000 cells/µL in DF-10S for transplantation.

### Flow cytometry and bOECs characterization

Cells were characterized by flow cytometry 1 month after plating. The number of primary bOECs were adjusted to a density of 2x10^5^ cells/ml PBS/ 0.5% bovine serum albumin (BSA) solution. TruStain FcX™ PLUS (BioLegend) was added to block non-specific binding. Cell populations were identified using anti-p75 nerve growth factor receptor (p75, Abcam, ab8874, RRID:AB_306827) and rat anti-platelet-derived growth factor β (PDGFRβ, Abcam, ab91066, RRID:AB_10563302) primary antibodies. OECs were identified as P75 positive and PDGFRβ negative cells. Stromal cells were identified as PDGFRβ positive and P75 negative cells. P75 and PDGFRβ primary antibody were revealed with the anti-rabbit phycoerythrin fluorochrome-conjugated (PE, poly4064, BioLegend, 406408, RRID:AB_10643424) and the anti-rat Alexafluor 488 fluorochrome-conjugated antibodies (AF488, MRG2b-85, BioLegend, RRID:AB_2715913) respectively. Data were analyzed using FlowJo software (version 10.3; FlowJo LLC).

### Real-time quantitative polymerase chain reaction characterization

Quantitative reverse transcription polymerase chain reaction (qRT-PCR) experiments were performed to evaluate the mRNA expression level of cytokines, chemokines and inflammatory molecules in OBs without SCI and 7 days after SCI and in BV2 cell cultures (BV2 control, BV2 + bOECs H conditioned medium (CM) and BV2 + bOECs I CM) (Table 1). In addition, qRT-PCR experiments were performed to measure mRNA expression levels of axon growth inhibitory, axon growth permissive molecules and neurotrophic factors in primary bOEC cultures (Table 2). Total RNA from bOECs was extracted after dissociation in the same manner as for the cultures, using Tri-reagent (Sigma) and Nucleospin RNAII Kit (Macherey-Nagel) according to the manufacturer’s protocol. From each sample, 1.5 µg of total RNA was converted into single-stranded cDNA using the ImPromII reverse transcriptase kit (Promega) with random primers (0.5 µg/ml). Two ng of complementary DNA (cDNA) were amplified in the presence of 2X SYBR Green Mastermix (Applied Biosystems) containing preset concentrations of dNTPs, MgCl2 and forward and reverse primers using a QuantStudioTM 12KFlex (Applied Biosystems). Appropriate controls without reverse transcriptase or with H2O instead of cDNA were performed. Primers were designed using Primer Express software (Applied Biosystems) and validated by RT-PCR. The relative amount of cDNA in each sample was determined using the comparative quantification cycle (Cq) method and expressed as 2-DDCq. Three housekeeping genes were used to standardize the relative amount of cDNA: gapdh, b2m and actb. All experiments were performed in duplicate.

**Table 1:**
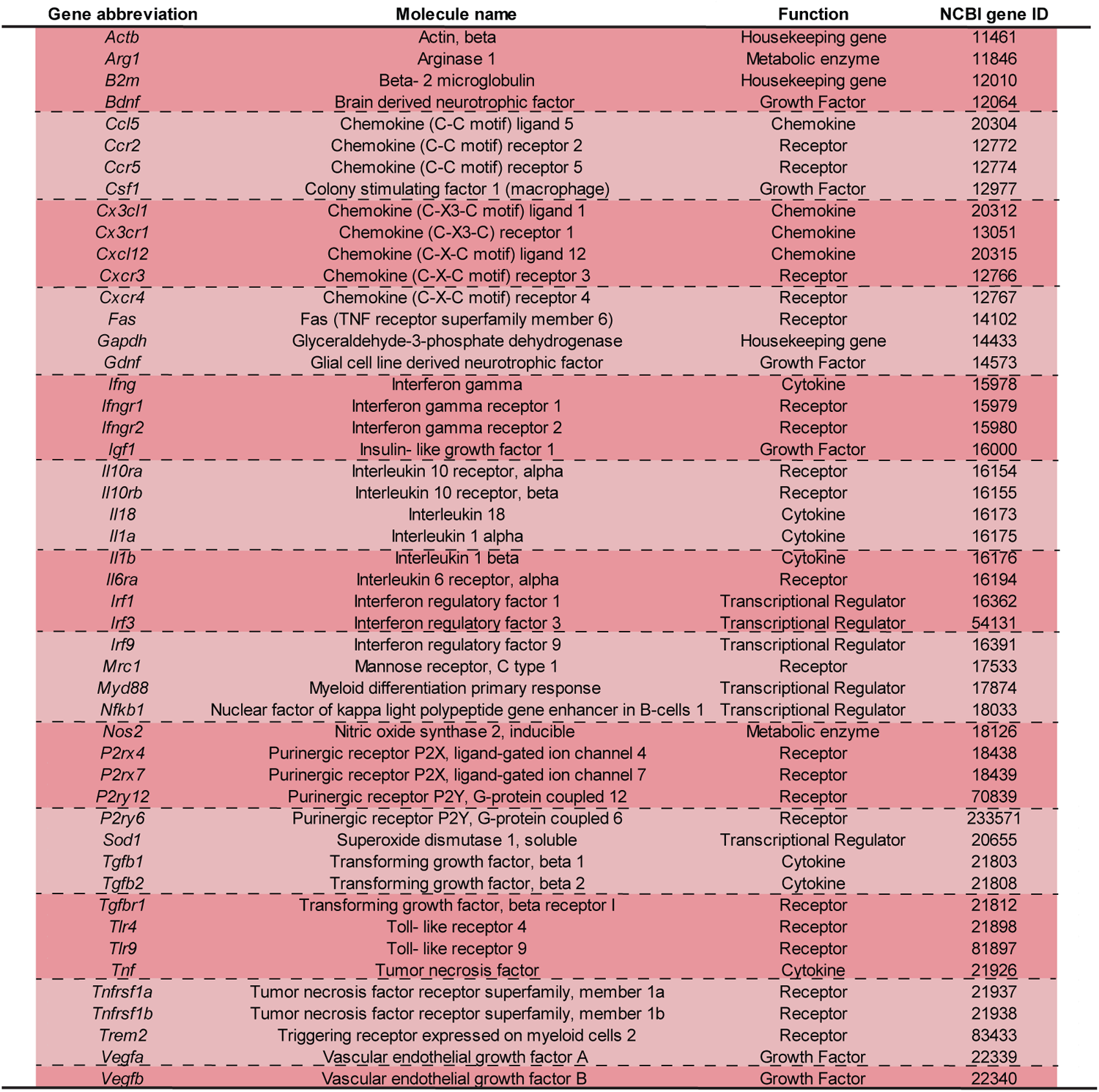
Table of genes for immunity and inflammatory molecules analyzed by qRT-PCR. This table lists the gene abbreviation, full name, function and NCBI gene ID of 49 immune and inflammatory molecules for which gene expression levels were analyzed in this study.

**Table 2:** Table of genes for axonal growth inhibitory and axonal growth permissive molecules analyzed by qRT-PCR. This table lists the gene abbreviation, full name, function and NCBI gene ID of 84 axon growth modulating molecules for which gene expression levels were analyzed in this study [5].

| Gene abbreviation | Molecule name | Function | NCBI gene ID |
| --- | --- | --- | --- |
| *Acan* | Aggrecan | ECM & Adhesion | 11595 |
| *Actb* | Actin beta | Housekeeping genes | 11461 |
| *Adam10* | ADAM metalloproteinase domain 10 | ECM & Adhesion | 11487 |
| *Bcan* | Brevican | ECM & Adhesion | 12032 |
| *Bdnf* | Brain derived neurotrophic factor | Growth Factor | 12064 |
| *B2m* | Beta-2-microglobulin | Housekeeping gene | 12010 |
| *Ccl2* | C-C Motif Chemokine Ligand 2 | Immunomodulatory | 20296 |
| *Cntf* | Ciliary Neurotrophic Factor | Axonal and synaptic plasticity | 12803 |
| *Col1a1* | Collagen type I alpha 1 chain | ECM & Adhesion | 12842 |
| *Col3a1* | Collagen type III alpha 1 chain | ECM & Adhesion | 12825 |
| *Col4a1* | Collagen type IV alpha 1 chain | ECM & Adhesion | 12826 |
| *Cspg4* | Chondroitin sulfate proteoglycan NG2 | ECM & Adhesion | 121021 |
| *Cspg5* | Neuroglycan C | ECM & Adhesion | 29873 |
| *Ctgf* | Connective tissue growth factor | Growth Factor | 14219 |
| *Dcc* | DCC netrin 1 receptor | Axonal and synaptic plasticity | 13176 |
| *Dcn* | Decorin | ECM & Adhesion | 13179 |
| *Efnb3* | Ephrin B3 | Axonal and synaptic plasticity | 13643 |
| *Eg-vegf* | Prokineticin 1 | Growth Factor | 246691 |
| *Epha4* | EPH receptor A4 | Axonal and synaptic plasticity | 13838 |
| *Ephb2* | EPH receptor B2 | ECM & Adhesion | 13844 |
| *Flt1* | Fms related tyrosine kinase 1 | Growth Factor | 14254 |
| *Flt4* | Fms related tyrosine kinase 4 | Growth Factor | 14257 |
| *Fn1* | Fibronectin 1 | ECM & Adhesion | 14268 |
| *Fgf1* | Fibroblast growth factor 1 | Growth Factor | 14164 |
| *Fgf2* | Fibroblast growth factor 2 | Growth Factor | 14173 |
| *Gapdh* | Glyceraldehyde-3-phosphate dehydrogenase | Housekeeping gene | 14433 |
| *Gdnf* | Glial derived neurotrophic factor | Growth Factor | 14573 |
| *Gpc1* | Glypican 1 | ECM & Adhesion | 14733 |
| *Gpc3* | Glypican 3 | ECM & Adhesion | 14734 |
| *Gpc5* | Glypican 5 | ECM & Adhesion | 103978 |
| *Gusb* | Glucuronidase beta | Housekeeping gene | 110006 |
| *Hsp90ab1* | Heat shock protein 90 alpha class B member 1 | Housekeeping gene | 15516 |
| *Ifng* | Interferon gamma | Immunomodulatory | 15978 |
| *Igf1* | Insulin like growth factor 1 | Growth Factor | 16000 |
| *Il1b* | Interleukine 1 beta | Immunomodulatory | 16176 |
| *Il6* | Interleukine 6 | Immunomodulatory | 16193 |
| *Kdr* | Kinase insert domain receptor | Growth Factor | 16542 |
| *Lamc1* | Laminin C 1 | ECM & Adhesion | 226519 |
| *Lglas1* | Galectin 1 | ECM & Adhesion | 16852 |
| *Matn2* | Matrilin 2 | ECM & Adhesion | 17181 |
| *Mcam* | Melanoma cell adhesion molecule | ECM & Adhesion | 84004 |
| *Mmp2* | Matrix metalloproteinase 2 | ECM & Adhesion | 17390 |
| *Mmp9* | Matrix metalloproteinase 9 | ECM & Adhesion | 17395 |
| *Met* | MET proto-oncogene, receptor tyrosine kinase | Growth Factor | 17295 |
| *Ncam1* | Neural cell adhesion molecule 1 | ECM & Adhesion | 17967 |
| *Ncan* | Neurocan | ECM & Adhesion | 13004 |
| *Neo1* | Neogenin 1 | ECM & Adhesion | 18007 |
| *Ntf3* | Neurotrophin 3 | Growth Factor | 18205 |
| *Nrp1* | Neuropilin 1 | Axonal and synaptic plasticity | 18186 |
| *Ntn1* | Netrin 1 | Axonal and synaptic plasticity | 18208 |
| *Pdgfa* | Platelet derived growth factor subunit a | Growth Factor | 18590 |
| *Pgf* | Placental growth factor | Growth Factor | 18654 |
| *Plxna1* | Plexin A1 | Axonal and synaptic plasticity | 18844 |
| *Plxnb1* | Plexin B1 | Axonal and synaptic plasticity | 235611 |
| *Rgma* | Repulsive guidance molecule BMP co-receptor A | Axonal and synaptic plasticity | 244058 |
| *Rgmb* | Repulsive guidance molecule BMP co-receptor B | Axonal and synaptic plasticity | 68799 |
| *Robo1* | Roundabout guidance receptor 1 | Axonal and synaptic plasticity | 19876 |
| *Robo2* | Roundabout guidance receptor 2 | Axonal and synaptic plasticity | 268902 |
| *Sdc1* | Syndecan 1 | ECM & Adhesion | 20969 |
| *Sdc2* | Syndecan 2 | ECM & Adhesion | 15529 |
| *Sdc3* | Syndecan 3 | ECM & Adhesion | 20970 |
| *Sdc4* | Syndecan 4 | ECM & Adhesion | 20971 |
| *Sema3a* | Semaphorin 3A | Axonal and synaptic plasticity | 20346 |
| *Sema3f* | Semaphorin 3F | Axonal and synaptic plasticity | 20350 |
| *Serpinf1* | Serpin family F member 1 | ECM & Adhesion | 20317 |
| *Slit1* | Slit guidance ligand 1 | Axonal and synaptic plasticity | 20562 |
| *Slitrk1* | SLIT and NTRK like family member 1 | Axonal and synaptic plasticity | 76965 |
| *Tek* | TEK receptor tyrosine kinase | Growth Factor | 21687 |
| *Tgfa* | Transforming growth factor alpha | Growth Factor | 21802 |
| *Tgfb1* | Transforming growth factor beta 1 | Growth Factor | 21803 |
| *Tgfb2* | Transforming growth factor beta 2 | Growth Factor | 21808 |
| *Tgfb3* | Transforming growth factor beta 3 | Growth Factor | 21809 |
| *Thbs1* | Thrombospondin 1 | ECM & Adhesion | 21825 |
| *Thbs2* | Thrombospondin 2 | ECM & Adhesion | 21826 |
| *Timp1* | Tissue Inhibitor Of Metalloproteinases 1 | ECM & Adhesion | 21857 |
| *Timp2* | Tissue Inhibitor Of Metalloproteinases 2 | ECM & Adhesion | 21858 |
| *Tnc* | Tenascin C | Axonal and synaptic plasticity | 21923 |
| *Tnf* | Tumoral necrosis factor | Immunomodulatory | 21926 |
| *Tnr* | Tenascin R | Axonal and synaptic plasticity | 21960 |
| *Vcan* | Versican | ECM & Adhesion | 13003 |
| *Vegfa* | Vascular endothelial growth factor A | Growth Factor | 22339 |
| *Vegfb* | Vascular endothelial growth factor B | Growth Factor | 22340 |
| *Ve-cad* | Vascular endothelial cadherin | ECM & Adhesion | 12562 |
| *Xylt1* | Xylosyltransferase 1 | ECM & Adhesion | 233781 |

### Surgical procedure and cell transplantation

SCI was performed at the T9-T10 level as previously described [19]. After dorsal incision and laminectomy of T7 vertebrae, incomplete spinal cord transection was performed to preserve the lower part of the ventral horns while completely damaging the central part and the dorsal horns of the spinal cord. The SCI was performed with a 25-gauge needle: cells were transplanted immediately after creating the lesion. bOECs were transplanted using a micromanipulator arm attached to a stereotactic frame (World Precision Instrument). Thus, bOECs were injected using a 1 mm sterile glass capillary needle attached to the micromanipulator. Injections were made at a depth of 1 mm, 1.5 mm from the midline, 5 mm rostral and caudal to the lesion site. Two injections of 2 µL containing 25,000 cells/µL of bOECs were delivered. Each injection was carefully administered within 1 minute. After surgery, the back muscles and skin were sutured and mice were checked daily, none of which showed skin lesions, infection or autophagy throughout the study.

### BDA injection

For axonal growth analysis, mice received an injection of 2 µL biotinylated dextran amine (BDA, 0.2 g/ml, Thermo Fisher Scientific, D1956, Waltham, MA) diluted in PBS at the cervical level two days prior to euthanasia. BDA was injected using a 1 mm sterile glass capillary needle attached to the micromanipulator arm mounted on a stereotactic frame (World Precision Instrument). Injections were made at a depth of 1 mm, 1.5 mm from the midline. The injection was carefully delivered within 1 minute. After injection, the muscles and skin were sutured.

### bOECs survival

The survival of LUX+/-bOECs was analyzed by bioluminescence using an *in vivo* XTREM 4XP spectrum cooled charge-coupled device optical macroscopic imaging system (Brucker). With intraperitoneal injections of D-luciferin (0.3 mg/g body weight, XenoLightTM, Perkin Elmer), LUX+/-bOECs were monitored for three weeks and photon counts were measured as previously described using Brucker molecular imaging software. [24].

In addition to the bioluminescence experiments, the survival of bOECs was investigated using bOECs extracted from Rosa26-tdTomato mice. After transplantation into wild type mice, the intensity of Rosa26-tdTomato bOECs was analyzed by immunohistochemistry (see below).

### Tissue preparation and sectioning

Thirty minutes before euthanasia, mice received analgesia (buprenorphine 0.17 mg/kg). Mice were then deeply anaesthetized with 5% isoflurane and given an intraperitoneal injection of sodium pentobarbital (100 mg/kg, euthoxin). Transcardial injections were made with cold PBS followed by cold 4% paraformaldehyde (PFA, Sigma Aldrich, 252549) in PBS. Dissected spinal cord and brain were post-fixed in 4% PFA overnight and cryoprotected in 30% sucrose (Life Technologies, Carlsbad, CA) for at least 48 hours. After embedding in Tissue-Tek OCT compound (Sakura, Tokyo, Japan), spinal cords were sectioned sagittally at 20 µm and brains sagittally at 30 µm. Sections were collected on slides and stored at -20°C until further use.

### Immunohistochemistry

Identification of different cells and structures in spinal cord and brain was performed using primary antibodies diluted in PBS blocking solution containing 10% normal donkey serum (Jackson ImmunoResearch, Cambridge, UK) and 0.3% Triton-X100 (Sigma-Aldrich) to block non-specific fixation. Antibody incubation was performed overnight at room temperature in a humidified chamber. The following primary antibodies were used: Rabbit anti-platelet-derived growth factor β (PDGFRβ, Abcam, ab32570, RRID:AB_777165), mouse anti-glial fibrillary acidic protein (GFAP Cy3-conjugated Sigma-Aldrich, C9205, RRID:AB_476889), rabbit anti-ionized calcium-binding adapter molecule 1 (Iba1, Wako, 019-19741, Osaka, Japan, RRID: AB_839504), rat anti-myelin basic protein (MBP, Millipore, MAB386, RRID:AB_94975), rabbit anti-laminin (Abcam, ab11575, RRID: AB_298179), goat anti-mMMR (CD206, R&D systems, AF2535, RRID:AB_2063012), rat anti-B-lymphocyte activation antigen B7-2 (CD86, Invitrogen, MA1-10299, RRID:AB_11152324), guinea pig anti-vesicular glutamate transporter 1 (vGlut1, Millipore, AB5905, RRID: AB_2301751) and Wisteria floribunda agglutinin biotin-conjugated lectin (WFA, Sigma, L1516, RRID: AB_2620171).

After washing, antibody staining was detected with species-specific fluorescence-conjugated secondary antibodies (Jackson ImmunoResearch). WFA antibody and BDA staining were visualized with Cy3-conjugated streptavidin (Jackson ImmunoResearch). Sections were counterstained with 4′,6-diamidino-2-phénylindole (DAPI; 1 μg/mL; Sigma-Aldrich) and cover slipped with Vectashield mounting medium (Vector Labs, Burlingame, UK).

### Image acquisition analysis

The representative images for quantification of immunohistochemically stained areas for sagittal sections were taken using the Zeiss Apotome2 microscope setup and the Leica THUNDER Imager Tissue 3D microscope setup. For spinal cord analysis, the epicenter of the lesion was imaged using a 10x microscope objective. For brain analysis, bulb images were taken using 20x microscope objective. Image processing and assembly was performed using ImageJ software.

### Quantification of immunohistochemically stained areas

Three to five sections per animal were examined to determine the epicenter of the lesion. An image of the epicenter was then taken and analyzed. On these sagittal sections, GFAP negative (-), PDGFrβ positive (+), MBP- and Laminin+ areas were measured. NeuN+, WFA+, Iba1+, CD86+, CD206+, Iba1+/IL1β+, Iba1+/IL4+ cells and BDA+/vGlut1+ axons were counted after standardization of the area considered with a rectangle of 1000 µm x 1000 µm.

### Primary OB conditioned-medium preparation

Primary OB cultures from uninjured and injured mice were prepared as described above. Four weeks after plating, the cultures were washed twice with PBS. The medium was then changed and CM was obtained 4 days later. The CM was placed in Amicon Ultra-15 3K ultrafiltration tubes (Millipore) and centrifuged at 5000g for 1 h at room temperature to concentrate it. From an initial volume of 6 mL, approximately 1 mL of concentrated CM was obtained by ultrafiltration. The CM was immediately stored at -80°C for preservation.

### BV2 Cell line cultures

The BV2 cell line was obtained from ATCC (EOC CRL-2467TM). Cells were first thawed gently in a water bath at 37°C. Cells were then cultured in T25 flasks containing DMEM high glucose (GIBCO), 20% fetal bovine serum (FBS), 0.5% penicillin/streptomycin for the first 48 hours. After this initial culture, a first passage was performed. To pass cells, the medium was removed and cells were washed twice with PBS. After adding 5 mL of trypsin EDTA, the flasks were placed in the incubator for 10 minutes. To inhibit trypsin, culture medium containing FBS was added. After centrifugation at 500g for 5 minutes, cells were resuspended in 1 mL of culture medium. The culture medium then consisted of DMEM glutaMAX high glucose (GIBCO), 10% FBS and 0.5% penicillin/streptomycin. One week after plating, cultures were passed, re-plated and divided into three experimental groups: 1 - BV2 control, 2 - BV2 + bOECs H CM and 3 - BV2 + bOECs I CM. Cells from these 3 groups were cultured for 3 days. Cultures were then trypsinized and immediately stored at -80°C for preservation for qRT-PCR analysis.

### Locotronic test: Foot misplacement apparatus

Experiments were performed as previously described by Chort et al. (Intellibio, Nancy, France) [25]. The apparatus consists of a flat ladder on which the animal can move from the start zone to the arrival zone. On both sides of the ladder, infrared sensors allow the visualization and recording of the animal’s movements. The location and precise length of time of all errors are recorded, in distinguishing errors from forelimb, hindlimb, and tail. Based on all recorded data, the number of hindlimb errors, total hindlimb error time and total crossing time were provided by the software and compared between groups of animals (Fig. 4B).

All mice were pre-trained on the ladder, three times 1 week prior to injury. Experiments were performed three times 14, 28 and 56 days after SCI for the bOECs transplanted groups.

### Hargreaves

A Hargreaves plantar test (Ugo basile®) was performed as previously described by [26]. Briefly, after isolation for 5 min, a radiant heat source was applied to the plantar surface of the hindpaw. The time between the onset of the thermal radiation stimulus and the onset of paw withdrawal was measured by computer. The time of paw withdrawal was defined as the hindpaw withdrawal latency. Experiments were performed 14, 28 and 56 days after SCI for the bOECs transplanted groups.

### CatWalk XT automated gait analysis

Motor coordination was performed and analyzed using the CatWalk XT automated system (Noldus; RRID:SCR_004074) [27]. In a dark room, mice were placed on the walkway and allowed to walk from one end to the other. The floor of the walkway consisted of a glass surface that transformed footprints of mice into light reflections in the form of illuminated footprints. The light reflection was recorded by a camera positioned under the walkway, the footprint image was converted into a digital signature and processed using CatWalk XT 9.1 software with a minimum threshold set at 80 (a.u. ranging from 0 to 225). After identification and labelling of footprints, data on static and dynamic gait parameters were generated from 3 consecutive trials (Fig. 4L and O). The average data of these parameters were analyzed and compared between experimental groups. If mice stopped halfway across or turned around during the run, the recording was stopped and the run was restarted to obtain 3 good crossings from one end to the other. Experiments were performed 56 days after SCI.

### Statistical analysis

All data are presented as average ± standard deviation (SD). All statistical analyses were performed using the GraphPad Prism program, version for windows (GraphPad Software; 8.0.1.244).

In the figure comparing two groups, the comparison of average was performed using the two-way Mann-Whitney test. In the figure comparing three groups, comparison of medians was performed using the two-way Kruskal-Wallis test followed by Dunn’s post-test. Statistical analysis in each assay is detailed in the figure legends. A p-value < 0.05 was considered statistically significant.

## RESULTS

### SCI alters gene expression in OBs tissue and primary bOEC cultures

To investigate the effects of SCI on distant tissues that could be used for autograft transplantation, we assessed the impact of SCI, 7 days after the lesion, on OB tissue and primary OB cultures by immunohistology, qRT-PCR and flow cytometry experiments (Fig. 2). A chronology of microglial cell reactivity in the OB was performed using immunohistology (Fig. 2A-D). Quantification of the total number of Iba1+ cells (Fig. 2A and B), the total number of double positive Iba1+/CD86+ cells (Fig. 2A and C) and the percentage of CD86+/Iba1+ cells among the Iba1+ cells (Fig. 2A and D) were assessed without SCI and at 3, 6, 12, 24, 48 hours and 7 days after injury. Our results show that the total number of Iba1+ cells does not change over time after SCI compared to the uninjured condition (Fig. 2B). On the contrary, the total number of double positive Iba1+/CD86+ cell is increased at 2 and 7 days after SCI compared to uninjured animals (Fig. 2C). Finally, our analyses indicate that the percentage of CD86+/Iba1+ cells among the Iba1+ cells is increased at 3 h after SCI and decreases to the control (No SCI) level at 12 h after SCI. There was a second rise of CD86+/Iba1+ cells 7 days after SCI in comparison to uninjured and to 12 h after SCI conditions (Fig. 2D).

**Figure 2:**
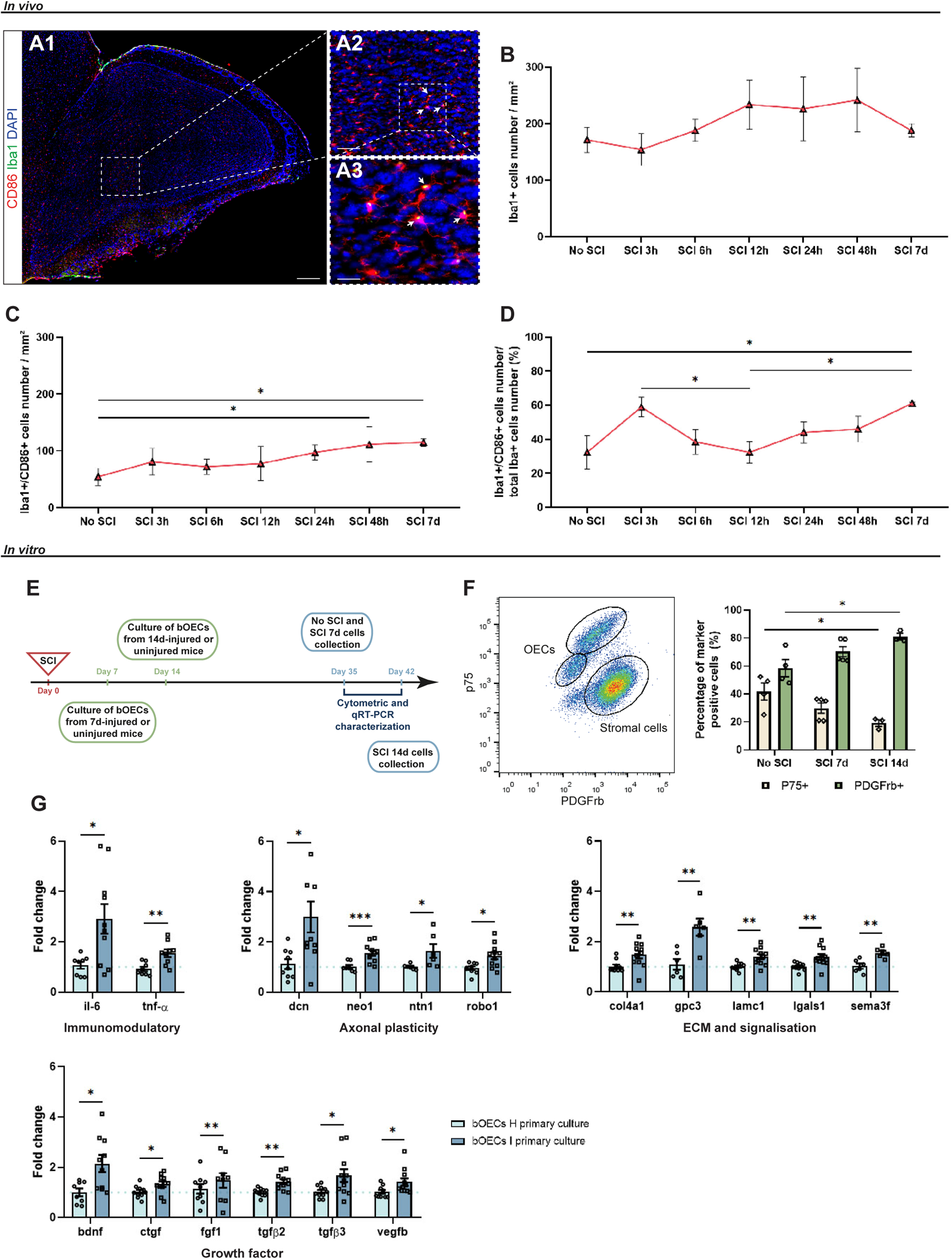
SCI alters cellular reactivity in olfactory bulbs and gene expression in cells in primary bOEC cultures. **A**. Representative sagittal section images of the granular cell layer in the OB. Sections were stained with anti-Iba1 and anti-CD86 antibodies. Microglial cell reactivity was analyzed by expression of Iba1 and CD86 markers. Arrows show Iba1+/CD86+ microglial cells. Scale bar: A1=250 µm, A2=100 µm and A3=20 µm. **B-D**. Chronology of microglial cell reactivity in the OB. Expression of cells markers Iba1 (**B**), CD86 (**C**) and the percentage of Iba1+/CD86+ cells (**D**) were measured without SCI and 3, 6, 12, 24, 48 hours and 7 days after SCI. **E**. Experimental design for bOEC H and bOEC I cultures characterization after SCI. Mice received SCI on day 0. Seven days after SCI, OB were collected from uninjured and 7d injured mice to establish bOEC H (No SCI) and 7d bOEC I (SCI 7d) primary cultures, respectively. Fourteen days after SCI, OB were collected from 14d injured mice to realize 14d bOEC I (SCI 14d) primary culture. At day 35, No SCI and SCI 7d primary cultures were characterized by cytometry and qRT-PCR. Characterization of SCI 14d primary culture was performed on day 42. **F**. Quantification of the percentage of p75+ OECs and PDGFRβ+ fibroblasts in bOEC cultures without SCI, 7 days and 14 days after SCI by cytometric analyses. **G**. Histograms of bOECs H and bOECs I mRNA expression of immunomodulatory, axonal plasticity, ECM/signaling and growth factor genes. Dashed lines correspond to mRNA expression from bOEC H cultures used as control. Quantifications are expressed as average ± SD. N=5 (**B-D**), 3 (**F**) and 8 (**G**) animals per group. Statistical evaluations were based on Mann-Whitney (**F** and **G**) and Kruskal-Wallis (**B-D**) tests. * = *P*< 0.05, ** = *P*< 0.01 and *** = *P*< 0.001.

Therefore, the effect of SCI on bOEC primary cultures was investigated (Fig. 2E-G). First, the composition of bOEC primary cultures was measured by flow cytometry. These analyses show that SCI modifies the ratio between OECs (p75-positive cells) and fibroblasts (PDGFrβ-positive cells) when cells are plated 14 days after SCI, but not at 7 days after SCI (Fig. 2F). qRT-PCR experiments were then performed on bOEC primary cultures 7 days after SCI and compared with bOEC primary cultures obtained from uninjured animals (Fig. 2G). Results show that SCI modulates the expression of genes involved in extracellular matrix (ECM) production and axonal plasticity processes, such as *sema3f*, *neo1*, *ntn1* or *lgals1*. In addition, SCI changes the expression of immunomodulatory genes (*il6* and *tnf-a*) and growth factor genes (e.g. *bdnf* or *vegfb*) in particular genes involved in scar-forming molecules such as *ctgf*, *fgf1*, *tgf*β*2* or *tgf*β*3* (Fig. 2G).

### The survival of bOECs I mice is identical to that of bOECs H after transplantation

To clarify whether the initial injury modulates the survival of bOECs after transplantation, bioluminescence and tdTomato positive bOEC transplantation experiments were performed up to 28 days after SCI (Fig. 3). Bioluminescence experiments show that in both transplanted groups (bOECs H and bOECs I) any surviving cells could be observed 21 days after SCI (Fig. 3A and B). These experiments also reveal that there is no difference between the two groups 7 and 14 days after SCI (Fig. 3A and B). To confirm these results, tdTomato positive bOECs were transplanted and immunohistological analyses were performed 14, 21 and 28 days after SCI (Fig. 3C and D). Measurements of tdTomato positive intensity show no difference between the two groups 14, 21 and 28 days after SCI (Fig. 3C and D).

**Figure 3:**
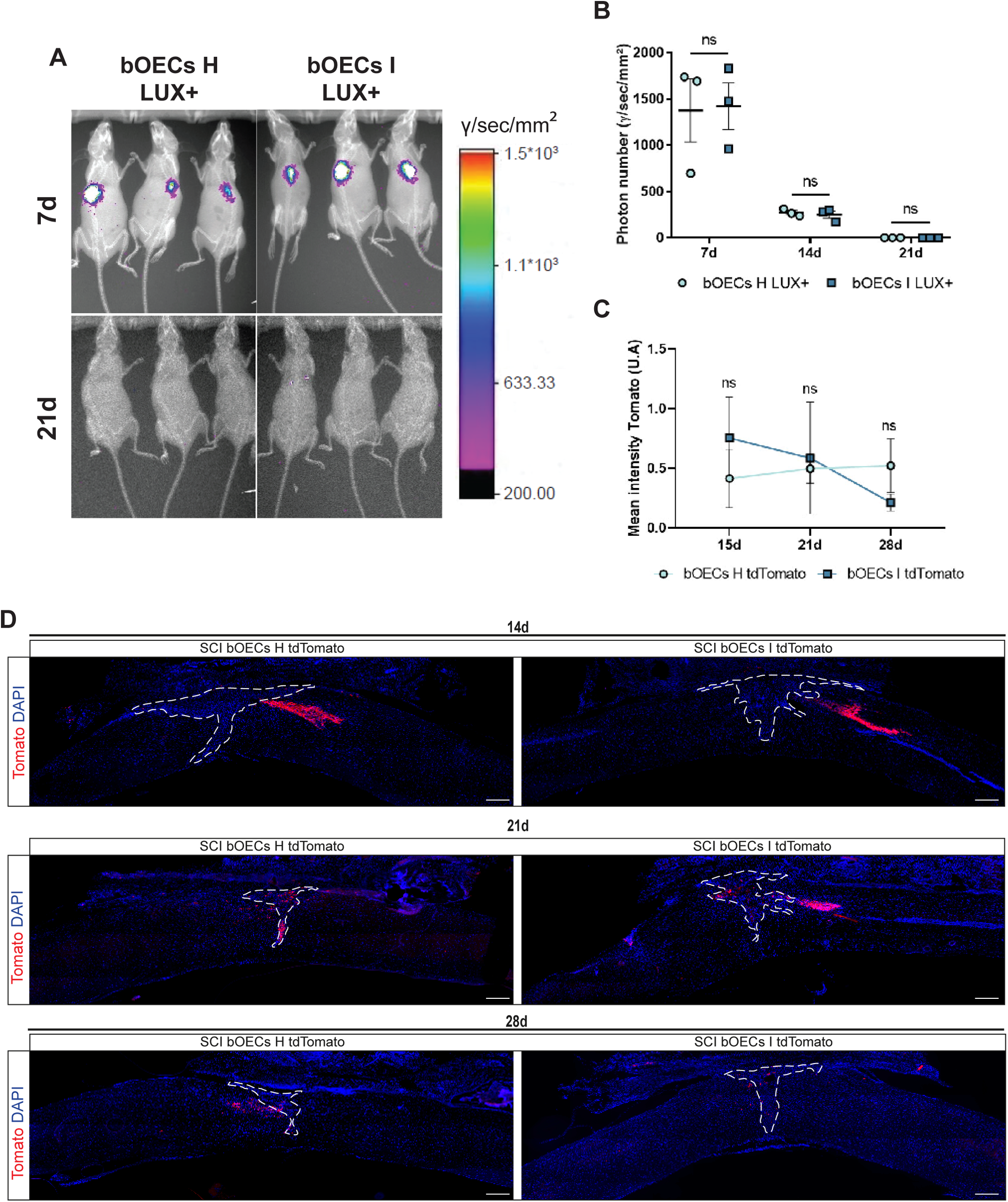
The survival of bOECs from injured mice is identical to bOECs’s survival from healthy mice after transplantation. **A**. Representative images of LUX+/-bOEC H and LUX +/-bOEC I cultures transplanted into wild type mice 7 and 21 days after SCI and transplantation. **B**. Quantification of luciferase+ signals 7, 14 and 21 days after SCI. **C**. Quantification of mean fluorescence intensity of the bOECs tdTomato H and I, 14, 21 and 28 days after SCI. **D**. Representative images of sagittal spinal cord sections from SCI bOECs H tdTomato and SCI bOECs I tdTomato groups 14, 21 and 28 days after SCI. Dashed lines correspond to the injury site. Scale bar = 250 µm. Quantifications are expressed as average ± SD. N=3 (**B**) and 5 (**C**) animals per group. Statistical evaluations were based on Kruskal-Wallis tests.

### Therapeutic capacity induced by bOEC transplantation is less important when bOECs are obtained from SCI mice

To assess whether the changes expressed at the RNA level in primary cultures of bOECs affect the neuroregenerative capacity of these cells after transplantation in the context of SCI, we evaluated functional recovery (Fig. 4A). First, the Locotronic test (Fig. 4B) was performed in the SCI, SCI + bOECs H and SCI + bOECs I groups 14, 28 and 56 days after SCI (Fig. 4C-K). In addition, fine motor movements were assessed using a CatWalk test at 56 days post SCI (Fig. 4L-O). Our analyses show that when bOECs are obtained from injured mice (bOECs I), their transplantation does not improve the number of hindlimb errors (Fig. 4C, F and I) at 14, 28 and 56 days after SCI, and only slightly improves the other parameters. In fact, except for the total hindlimb error time 14 days after SCI (Fig. 4D) and the time needed to cross the corridor 28 days after SCI (Fig. 4H), Locotronic test reveals that bOECs I group is comparable to SCI group.

**Figure 4:**
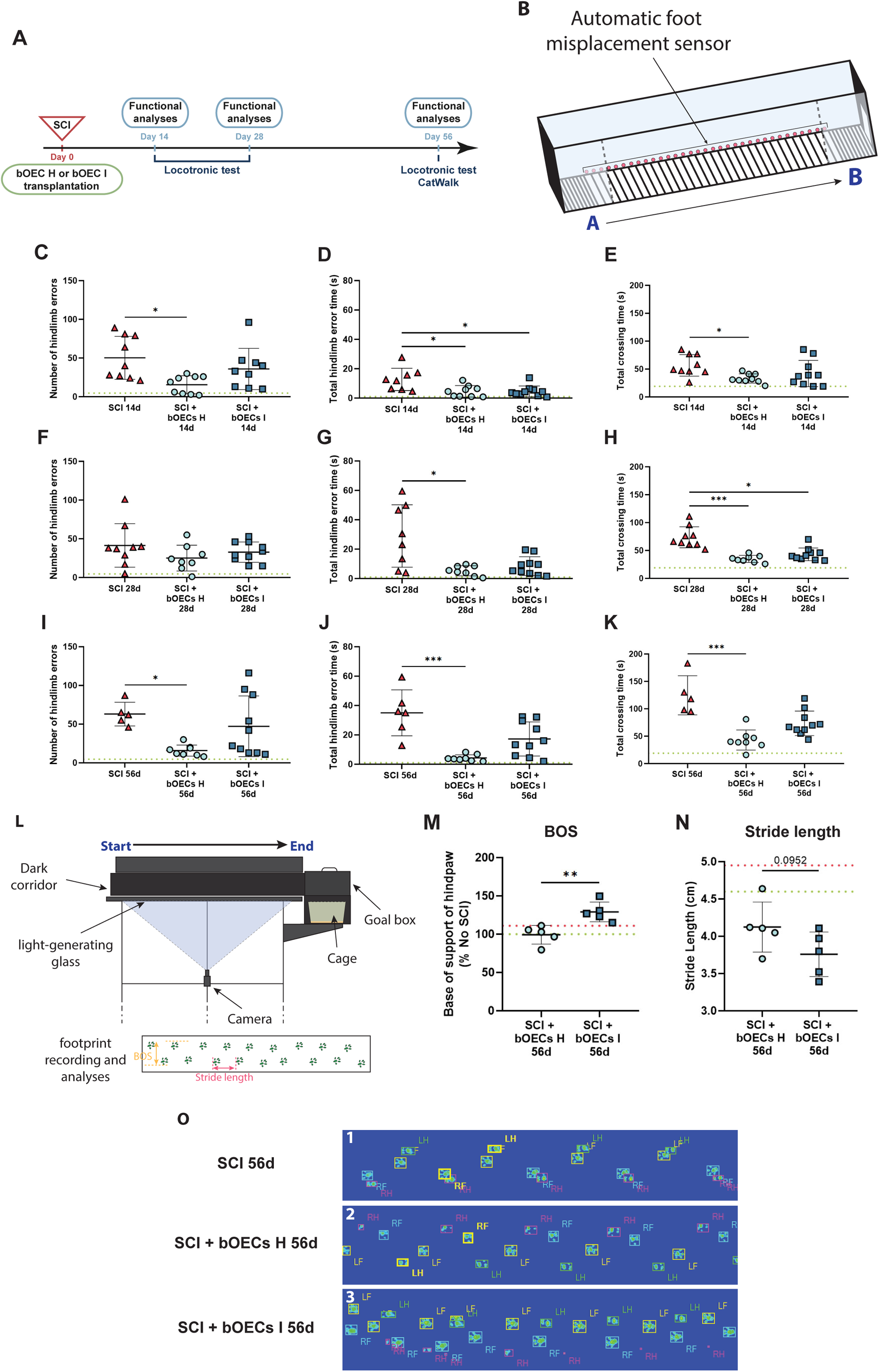
Transplantation of bOECs from injured mice resulted in less functional recovery than bOECs from healthy mice. **A**. Experimental design for experiments on motor behavior. Mice underwent SCI and bOEC H or I transplantation on day 0. Fourteen, twenty-eight and fifty-six days after SCI, functional analyses were performed. **B**. Schematic representation of the Locotronic test. **C**-**K**. Functional recovery was analyzed using Locotronic test 14, 28 and 56 days after SCI. Dashed lines correspond to the baseline parameters obtained during Locotronic test habituation (7 days before SCI). **C**, **F** and **I**. Quantification of the number of hindlimb errors. **D**, **G** and **J**. Quantification of the total hindlimb errors time. **E**, **H** and **K**. Quantification of the total crossing time. **L**. Schematic representation of the CatWalk test. **M** and **N**. CatWalk gait analysis was performed 56 days after SCI. Green and red dashed lines corresponds respectively to the baseline parameters obtained with CatWalk gait analysis during habituation (7 days before SCI) and to parameters obtained by SCI group 56 days after SCI. **M**. Measurement of the base of support (BOS). **N**. Measurement of the stride length. **O**. Example of photographs taken during the CatWalk gait for analyses. 1-SCI group, 2-SCI + bOECs H group, 3-SCI + bOECs I group. Quantifications are expressed as average ± SD. N=8-10 (**C**-**H**), 5-10 **(I**-**K**) and 5 (**M** and **N**) animals per group. Statistical evaluations were based on Kruskal-Wallis (**C**-**K**) and Mann-Whitney (**M** and **N**) tests. Quantifications are expressed as average ± SD. * = *P*< 0.05; ** = *P*< 0.01 and *** = *P*< 0.001.

To confirm these results, CatWalk experiments were performed 56 days after SCI. The results obtained show that the base of support (BOS) (Fig. 4M) is increased in bOECs I compared to bOECs H mice, also the stride length tends to be reduced in the bOECs I group compared to bOECs H (Fig. 4N).

The Locotronic test and CatWalk analyses indicate that the improvement in both static and dynamic parameters is lost in bOECs I group compared to the bOECs H group. To correlate these results with functional recovery, immunohistological analyses were performed 14, 28 and 56 days after SCI (Fig. 5). Spinal scar and demyelination were assessed by quantification of fibrotic (Fig. 5B, F, J and M) and glial scar (Fig. 5C, G, K and N) and quantification of demyelinated areas (Fig. 5D, H, L and O) over time. As previously described, bOEC H transplantation modulates the lesion scar by decreasing the fibrotic scar (PDGFrβ+ area) and demyelination (MBP-area) after SCI and by increasing the glial scar (GFAP-area) 14 and 56 days after SCI (Fig. 5A, B, C, D, I, J, K and L), with the same tendency without statistical difference 28 days after SCI (Fig. 5E-H) [28]. More interestingly, immunohistological results show that bOEC I transplantation does not modulate fibrotic and glial scars 14 (Fig. A-C), 28 (Fig. E-G) and 56 days (Fig. I-K) post SCI. Analysis of the demyelination process indicates that bOEC I transplantation reduces the MBP negative area only at 56 days post SCI, but not at 14 and 28 days post SCI (Fig. 5L, D and H respectively). The time course of SCI allows us to observe that at 14 and 28 days post SCI, the spinal cord scar is larger in animals that had received a bOEC I graft but at 56 days post SCI, the scar size tends to be identical to that of animals who had received bOECs H (Fig. M-O).

**Figure 5:**
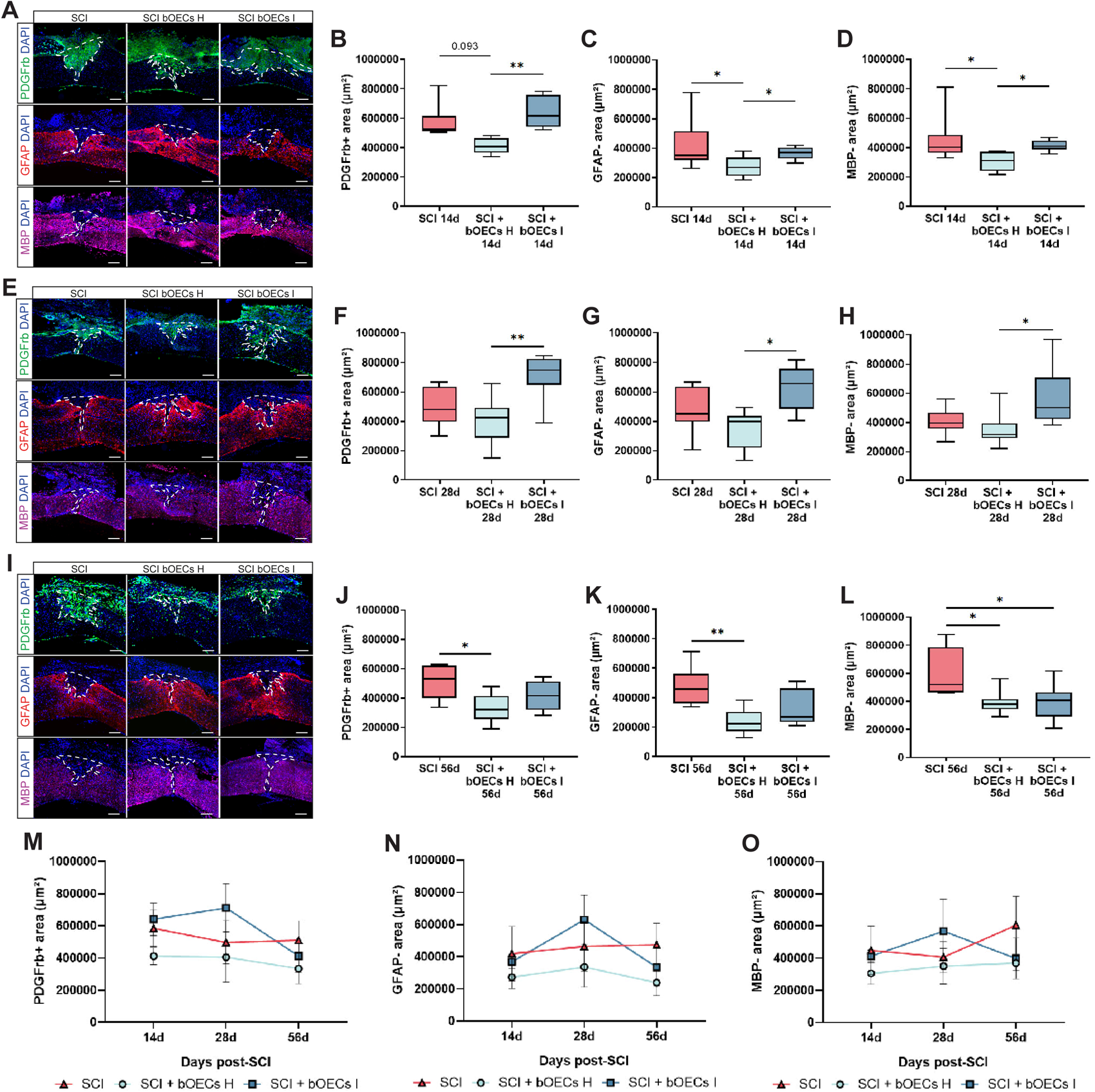
The tissue repair properties induced by transplanting bOECs from healthy mice are not achieved by transplanting bOECs from injured mice. Histological analyses of the spinal scar were performed 14, 28 and 56 days after SCI. **A, E** and **I**. Representative pictures of sagittal spinal cord sections of SCI, SCI bOECs H and SCI bOECs I groups 14 (**A**), 28 (**E**) and 56 (**I**) days after SCI. Sections were stained with anti-GFAP, anti-PDGFRβ and anti-MBP antibodies. **B, F** and **J**. Quantification of fibrosis areas (PDGFRβ+) 14 (**B)**, 28 (**F**) and 56 (**J**) days after SCI. **C, G** and **K**. Quantification of astrocyte-free areas (GFAP-) 14 (**C**), 28 (**G**) and 56 (**K**) days after SCI. **D, H** and **L**. Quantification of demyelinated/unmyelinated area (MBP-) 14 (**D**), 28 (**H**) and 56 (**L**) days after SCI. **M**-**O**. Representation of changes over time of the three analyses: PDGFRβ+ (**M**), GFAP-(**N**) and MBP-(**O**). **A, E** and **I**. Dashed lines correspond to the injury site. Scale bar = 200 µm. Quantifications are expressed as average ± Min/Max. N=8-10 (**B, C, D, F, G** and **H**), 5-10 (**J-L**) animals per group. Statistical evaluations were based on Kruskal-Wallis tests. Quantifications are expressed as average ± SD. * = *P*< 0.05 and ** = *P*< 0.01.

Collectively, our results demonstrate that the initial trauma induced prior to primary culture of bOECs alters their ability to promote functional recovery and tissue repair after SCI and transplantation.

### Transplantation of bOECs induces axonal regrowth, but this regrowth is associated with synaptogenesis and pain-related behavior only in mice that received bOECs from healthy mice

It has been previously described that cell transplantation, including transplantation of OECs, can induce allodynia after SCI [29]. To investigate this, the Hargreaves plantar test was performed 14, 28 and 56 days after SCI (Fig. 6A and B). In addition, hindpaw pressure was assessed using the CatWalk system 56 days after SCI (Fig. 6A, C and D). As previously described, bOEC H transplantation reduces hindpaw withdrawal time 14 days after SCI compared to SCI mice (Fig. 6B). Experiments also demonstrate that bOECs I transplanted animals had a hindpaw withdrawal time comparable to the SCI group, which was also increased compared to the bOECs H group 14 and 56 days after SCI (Fig. 6B). To confirm these results, the hindpaw pressure of the hindlimbs was measured 56 days after SCI. Measurements show that maximum and mean hindpaw pressures tend to decrease with transplantation of bOECs H compared to bOECs I (Fig. 6C and D), which may indicate pain-related behavior during locomotion in bOECs H mice.

**Figure 6:**
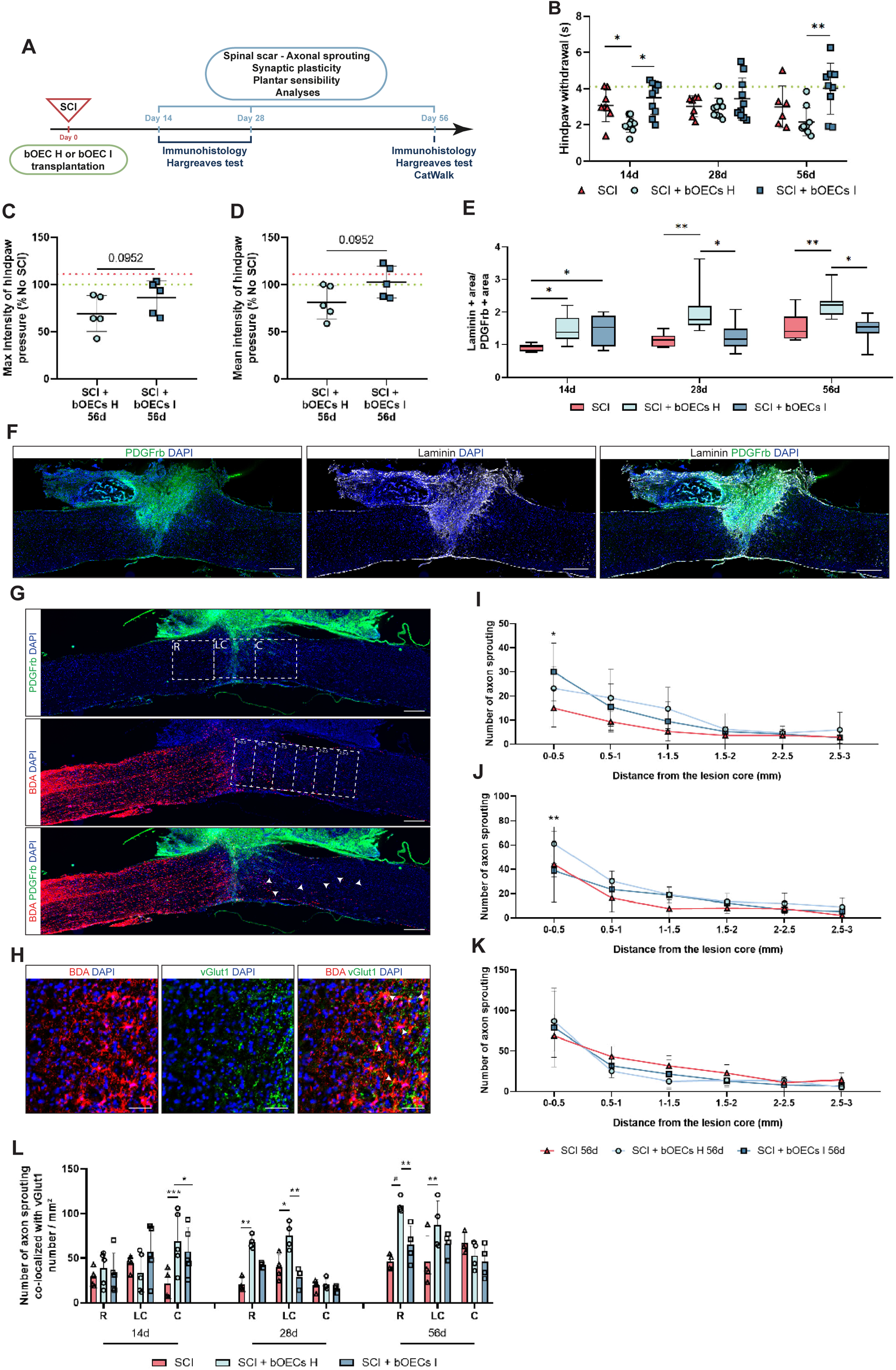
Transplantation of bOECs induces axonal regrowth, but this regrowth is associated with synaptogenesis and pain-related behavior only in mice that received bOECs from healthy mice. **A**. Experimental design for experiments on sensory behavior. Mice received SCI and bOEC H or I transplantation on day 0. Fourteen, twenty-eight and fifty-six days after SCI, functional analyses and immunohistology were performed. (**B**) Functional sensory recovery was analyzed 14, 28 and 56 days after SCI, using the Hargreaves test. Dashed lines correspond to the baseline results obtained during the Hargreaves test habituation (7 days before SCI). **C** and **D**. CatWalk gait analysis was performed. Quantification of hindpaw pressure measurements, maximum (**C**) and mean (**D**) hindpaw pressure measurements at 56 days after SCI. Green and red dashed lines correspond respectively to the baseline results obtained during the CatWalk gait test habituation (7 days before SCI) and to the results obtained by the SCI group 56 days after SCI. In addition, (**E-L**) histological analyses were performed 14, 28 and 56 days after SCI. **E-F**. Laminin expression was analyzed as a support for axonal growth. Quantification of laminin+ area/ PDGFRβ+ area ratio (**E**) 14, 28 and 56 days after SCI. **F**. Representative images of sagittal spinal cord sections from SCI bOECs I group 14 days after SCI, showing the laminin area and corresponding PDGFRβ area. Sections were stained with anti-laminin antibody and anti-PDGFRβ antibodies. Scale bar = 500 µm. **G**-**H**. Representative images of sagittal spinal cord sections from SCI bOECs I group. **G**. The square sections correspond to the rostral (R), lesion core (LC) and caudal (C) areas, defined for quantification of BDA+/vGlut1+ axon sprouting. The rectangular sections correspond to the areas defined for the quantification of axon sprouting as a function of distance from the lesion core. Arrows indicate axon sprouting (**G**) and co-localized BDA+/vGlut1+ axon (**H**). Sections were stained with (**G**) biotinylated BDA and anti-PDGFRβ antibody and (**H**) biotinylated BDA and anti-vGlut1 antibody. Scale bar = 500 µm (**G**) and 100 µm (**H**). **I-K**. The number of sprouting axons as a function of distance from the lesion core was counted 14, 28 and 56 days after SCI. **L**. Sprouting axons co-localized with synaptic vGlut1+ were counted 14, 28 and 56 days after SCI. Quantifications are expressed as average ± SD (**B, C, D, I, J, K,** and **L**) and average ± Min/Max (**E**). N=6-10 (**B** and **E**) and 5 (**C, D, I, J, K,** and **L**) animals per group. Statistical evaluations were based on Kruskal-Wallis (**B, E, I, J, K,** and **L**) and Mann-Whitney (**C** and **D**) tests. * = *P*< 0.05; ** = *P*< 0.01 and *** = *P*< 0.001.

Previous studies have described that pain-related behavior can be associated with axonal regrowth and formation of new synapses [30]. The composition of ECM plays a crucial role in axonal regrowth or axonal inhibitory processes as well as synaptogenesis. Therefore, the presence of laminin (Fig. 6A, E and F), an ECM molecule permissive that promotes axonal growth, was examined. As the size of the laminin area is related to the size of the lesion core (PDGFβ+ area), we measured the ratio of laminin+ area to PDGFβ+ area. It appears that the relative size of the laminin+ area is increased in both transplanted groups compared to SCI group 14 days after SCI and only in the bOECs H group compared to the SCI and bOECs I groups 28 and 56 days after SCI (Fig. 6E and F).

To investigate axonal regrowth and the formation of new synapses, BDA anterograde tracing experiments (Fig. 6A, G, I, J and K) and colocalization analysis of BDA and VGlut1 staining (Fig. 6A H and L) were performed. As previously described [31], measurement of the number of BDA-positive axons present in the lesion core shows that bOEC transplantation induces modest axonal regrowth after SCI compared to the SCI group (Fig. 6I-K). Indeed, our results indicate that bOECs I mice show an increase in the number of BDA+ fibers in the lesion core 14 days after SCI, whereas bOECs H induces it 28 days after SCI (Fig. 6I and J respectively). Fifty-six days after SCI, no difference is observed between groups (Fig. 6K). In addition, colocalization analysis of BDA and VGlut1 staining were performed 14, 28 and 56 days after SCI (Fig. 6A, H and L). This analysis shows that bOEC H transplantation increases synaptogenesis at the rostral part of the spinal cord compared to SCI groups and at the lesion core 28 days after SCI compared to the SCI group and bOECs I group (Fig. 6L). Fifty-six days after SCI, bOEC H transplantation also enhances synaptogenesis at the rostral part of the spinal cord compared to the SCI group and at the lesion core compared to bOECs I group and to the SCI group (Fig. 6L). In contrast, bOEC I transplantation did not modulate synaptogenesis, except at 14 days after SCI, where bOEC I transplantation increased it at the caudal part of the spinal cord compared to the SCI group (Fig. 6L). These results tend to demonstrate that bOEC H transplantation induces a more supportive microenvironment for axonal regrowth.

Taken together, our results indicate that bOEC H transplantation modifies the lesion scar ECM, which enhances axonal regrowth and synaptogenesis, and that bOEC I transplantation plays a weaker role on these parameters.

### Transplantation of bOECs from injured animals fails to improve neuronal survival

Neuronal survival was assessed in the three groups 14, 28 and 56 days after SCI (Fig. 7A, B, C, D, G and J). NeuN+ cells were counted at the rostral, caudal and lesion core of the spinal cord. As described in the literature, bOEC H transplantation increases neuronal survival [32]. Indeed, our results show that 14 days after SCI, neuronal survival is increased in bOECs H animals at the rostral part of the spinal cord compared to SCI group (Fig. 7D). Twenty-eight days after SCI, bOEC H transplantation increases the number of NeuN+ cells in both rostral and caudal parts of the spinal cord compared to SCI animals (Fig. 7G). Finally, 56 days after SCI, results show no difference between the groups (Fig. 7J). These analyses demonstrate that bOEC I transplantation does not improve neuronal survival 14, 28 and 56 days after SCI.

**Figure 7:**
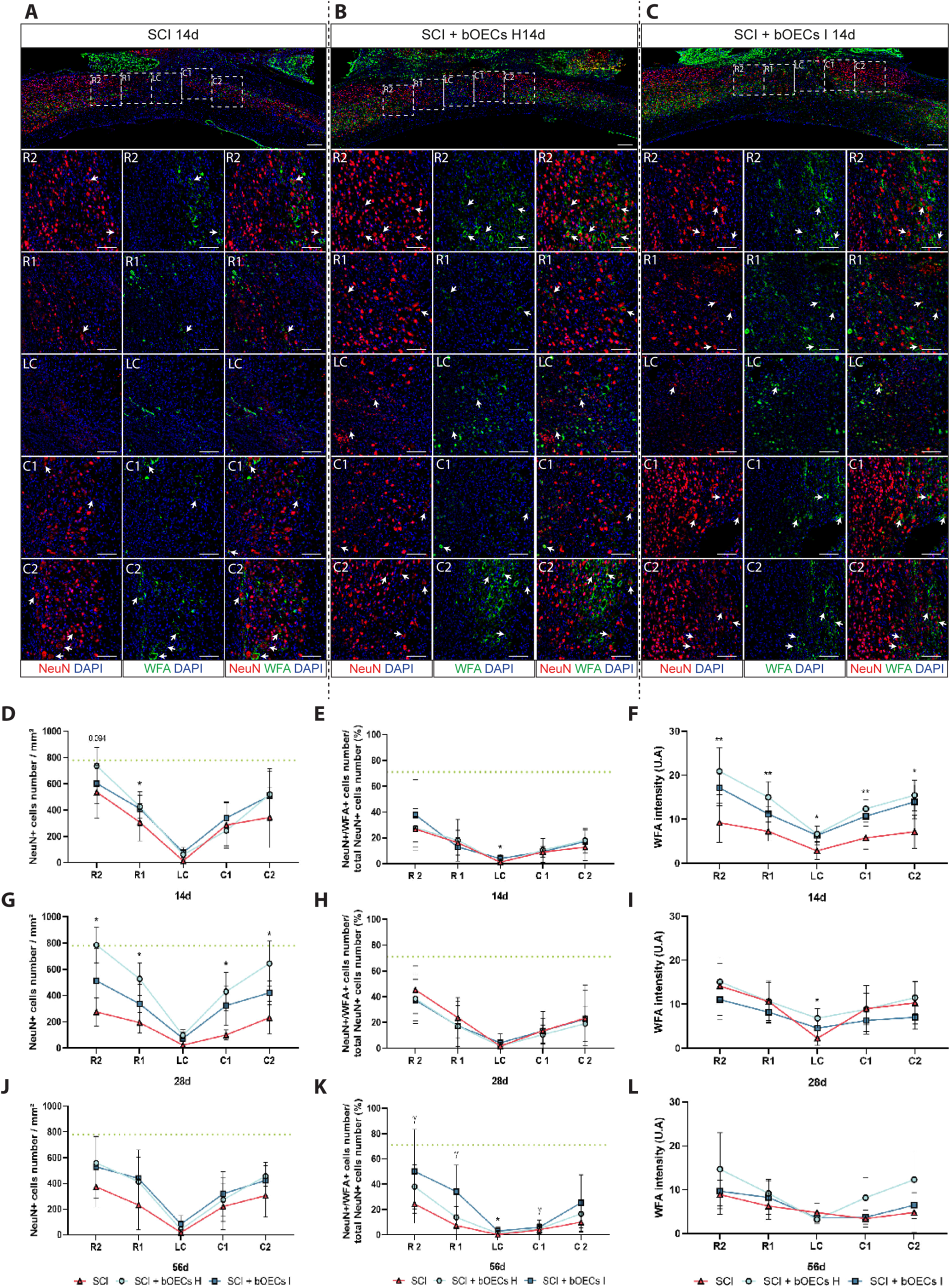
Transplantation of bOECs from healthy and injured mice affects neuronal survival and perineural network structure differently. Histological analyses of neuronal density and stability. **A**-**C**. Representative images of sagittal spinal cord sections from SCI (**A**), SCI bOECs H (**B**) and SCI bOECs I (**C**) groups 14 days after SCI. Sections were stained with anti-NeuN antibody and biotinylated WFA. The square sections correspond to the rostral 1 and 2 (R1 and R2), lesion core (LC) and caudal 1 and 2 (C1 and C2) areas, defined for the quantification of NeuN+ neurons and WFA+ perineural net. Arrows indicate co-localized NeuN and WFA staining. Scale bar = 250 µm (sagittal spinal cord sections) and 100 µm (R2, R1, LC, C1 and C2 squares analyzed). **D**, **G** and **J**. Quantification of the neuronal density 14 (**D**), 28 (**G**) and 56 (**J**) days after SCI. **E**, **F**, **H**, **I**, **K** and **L**. Analysis of the perineural net. **E**, **H** and **K**. Quantification of the number of NeuN+ neurons surrounded by WFA 14 (**E**), 28 (**H**) and 56 (**K**) days after SCI. **F**, **I** and **L**. Measurement of WFA intensity 14 (**F**), 28 (**I**) and 56 (**L**) days after SCI. Dashed lines correspond to the baseline parameters obtained in the No SCI group. Quantifications are expressed as average ± SD. N=5-8 (**D**-**F**), 5-10 (**G**-**I**) and 5 (**J**-**L**) animals per group. Statistical analysis was based on Kruskal-Wallis test. * = *P*< 0.05 and ** = *P*< 0.01.

We then measured the number of NeuN+/WFA+ cells at rostral and caudal parts and at the lesion core of the spinal cord 14, 28 and 56 days after SCI. Fourteen days after SCI, the number of NeuN+/WFA+ cells is increased in the bOECs H group at the lesion core compared to the SCI group (Fig. 7E), with no difference between groups 28 days after SCI (Fig. 7H). Interestingly, at 56 days post SCI, our results indicate that the number of NeuN+/WFA+ cells is increased in bOECs H group compared to the SCI group at the lesion core, whereas the number of NeuN+/WFA+ cells is increased in the rostral and caudal parts of the spinal cord in bOECs I animals compared to the SCI group (Fig. 7K).

In addition to quantifying the number of NeuN+/WFA+ cells, WFA intensity was also assessed at 14, 28 and 56 days after SCI at the rostral and caudal parts and at the lesion core of the spinal cord (Fig. 7F, I and L). Fourteen days after SCI, bOEC transplantation improves WFA intensity quantifications at rostral and caudal parts and at the lesion core with a tendency for bOECs I group compared to SCI group and with significant differences for bOECs H group (Fig. 7F). At 28 days after SCI, quantification of WFA intensity shows that bOEC H transplantation increases this parameter at the lesion core compared to the SCI group, with no difference between groups at 56 days after SCI (Fig. 7I and L respectively). At these time points, no difference is observed between SCI + bOECs I group and SCI group (Fig. 7I and L).

### Transplantation of bOECs modulates the inflammatory microenvironment after SCI with moderate effects when bOECs are obtained from injured mice

The immunomodulatory role of OECs after transplantation has recently been highlighted in several studies (Jiang et al., 2022). Therefore, we investigated immunomodulatory properties of bOEC H and bOEC I transplantation at 14, 28 and 56 days after SCI (Fig. 8). First, the number of Iba1+ cells present at rostral and caudal parts of the spinal cord and in the lesion core was measured. This analysis shows that there is no difference between the groups overtime (Fig. 8A). Macrophages and microglial cells can express anti-inflammatory or pro-inflammatory profiles with the respective expression of CD206 and CD86 markers. The ratio of CD86+/Iba1+ and CD206+/Iba1+ cells were assessed 14, 28 and 56 days after SCI at rostral and caudal parts of the spinal cord and in the lesion core (Fig. 8B-I). Fourteen days after SCI, bOEC H transplantation decreased the ratio of CD86+/Iba1+ cells at rostral part of the spinal cord and increased the ratio of CD206+/Iba1+ cells at rostral and caudal parts of the spinal cord compared to SCI animals. Transplantation of bOECs I tends to change the profile of microglia/macrophages in the same way at this time point, with no significant difference compared to the SCI group (Fig. 8D and E). Twenty-eight days after SCI, both bOECs transplanted groups modulate the microglia/macrophage profile (Fig. 8F and G).

**Figure 8:**
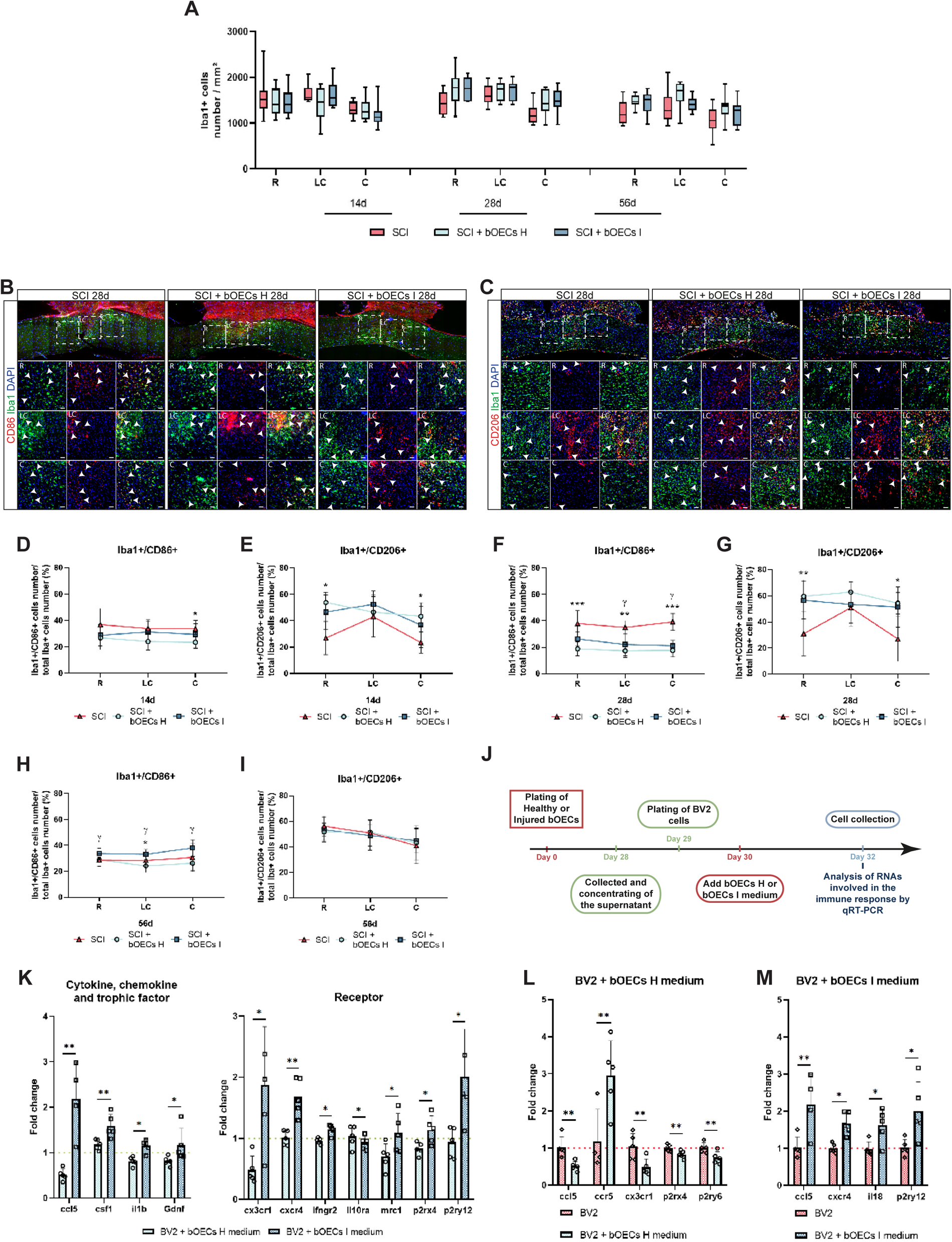
Transplantation of bOECs from healthy and injured mice alters microglia/macrophages polarization. **A-I**. Histological analysis of microglia/macrophage cell density and polarization. **A.** The number of Iba1 positive microglia/macrophage cells was measured 14, 28 and 56 days after SCI. **B** and **C**. Representative images of sagittal spinal cord sections from SCI, SCI bOECs H and SCI bOECs I groups 28 days after SCI. Sections were stained with anti-Iba1 and anti-CD86 antibodies (**B**) and anti-Iba1 and anti-CD206 antibodies (**C**). The square sections correspond to the rostral (R), lesion core (LC) and caudal (C) areas, defined for the quantification of Iba1+/CD86+ and Iba1+/CD206+ cells. Arrows indicate Iba1+/CD86+ microglial cells (**B**) and Iba1+/CD206+ microglial cells (**C**). Scale bar = 250 µm (sagittal spinal cord sections) and 100 µm (Squares analysed in R, LC and C). Microglia/macrophage polarization was analyzed by expression of pro-inflammatory marker CD86 (**D, F** and **H**) and anti-inflammatory marker CD206 (**E, G** and **I**) at 14 (**D** and **E**), 28 (**F** and **G**) and 56 (**H** and **I**) days post SCI. **J-M**. Induction of BV2 cells polarization *in vitro* by bOECs conditioned medium (CM). **J**. Experimental design, healthy or injured bOECs were plated (day 0). Four weeks after plating, bOECs H and bOECs I CM were collected and concentrated (day 28). BV2 cells were plated (day 29) and the next day (day 30), bOECs H or bOECs I concentrated CM was added to the BV2 cell cultures. Two days after (day 32), BV2 cells were removed from the plates and analyzed by qRT-PCR. **K-M**. Histogram of mRNA expression of BV2 cell control and BV2 cells with bOECs H CM or bOECs I CM of cytokine, chemokine, trophic factor and receptor genes. Dashed lines correspond to mRNA expression from control BV2 cells cultures. Quantifications are expressed as average ± SD (**D**, **E**, **F**, **G**, **H**, **I**, **K**, **L** and **M**) and average ± Min/Max (**A**). N=5-10 (**A**), 8-10 (**D-G**) and 5-10 (**H** and **I**) animals per group. N=5 BV2 cell cultures (**K, L** and **M**) per group. Statistical analyses were based on Kruskal-Wallis (**A, D, E, F, G, H** and **I**) and Mann-Whitney (**K, L** and **M**) tests. * and γ = *P*< 0.05; ** = *P*< 0.01 and *** = *P*< 0.001.

Indeed, the ratio of CD86+/Iba1+ cells is reduced at the lesion core and at the caudal part of the spinal cord in both transplanted groups compared to SCI animals and is also reduced at the rostral part of spinal cord in bOECs H group (Fig. 8F). The analysis also reveals that the ratio of CD206+/Iba1+ cells is increased in both transplanted groups at rostral and caudal parts of the spinal cord compared to the SCI group, with a significant difference only between the bOECs H and SCI groups (Fig. 8G). Interestingly, 56 days after SCI, bOEC I transplantation increased the ratio of CD86+/Iba1+ cells at rostral and caudal part and lesion core of the spinal cord compared to bOECs H animals (Fig. 8H). Furthermore, the ratio of CD86+/Iba1+ cells at the lesion core is also increased between bOECs I and SCI groups. No difference is observed in CD206+/Iba1+ ratio analyses between groups (Fig. 8L).

To complete the analysis of bOECs properties in modulating inflammation, we performed qRT-PCR analyses on BV2 cell cultures (microglial cell line) exposed to conditioned media obtained from bOEC H (BV2 + bOECs H medium) and bOEC I (BV2 + bOECs I medium) cultures (Fig. 8J). The results obtained show that chemoattractant (*ccl5*) or inflammation activator genes (e.g. *csf1* or *cx3cr1*) are upregulated in BV2 + bOECs I compared to BV2 + bOECs H (Fig. 8K). It also appears that pro-inflammatory genes (e.g. *il1b*, *cxcr4*, *ifngr2*) are upregulated in BV2 + bOECs I compared to BV2 + bOECs H and anti-inflammatory genes are downregulated (*il10ra*). The comparison between BV2 and BV2 + bOECs H shows that activated stage (*p2ry12* and *p2rx4*) and pro-inflammatory (*ccl5* and *cx3cr1*) genes are downregulated and pro-remyelinating (*ccr5*) gene is upregulated in BV2 + bOECs H (Fig. 8L). Whereas pro-inflammatory genes (e.g. *ccl4* or *cxcr4*) are upregulated in BV2 + bOECs I compared to BV2 (Fig. 8M).

Collectively, our results demonstrate that the initial injury modulates bOECs I CM, which in turn alters gene expression of BV2 cell lines *in vitro*. Interestingly, these effects found *in vitro* do not strongly modify the *in vivo* properties of bOECs after transplantation in the context of SCI.

## DISCUSSION

### Spinal cord injury and regeneration by bOEC transplantation

The main purpose of our study was to investigate SCI consequences on brain and peripheral tissues. In particular, on the impairment of cell therapeutic capacities that could be used in an autograft context. Cellular responses and tissue remodeling that occur after SCI are well described in spinal tissue but only few studies have investigated the cellular responses induced by the trauma in the whole body. SCI is characterized by neuronal, glial, microglial and peripheral immune cells activation. Immune cells induce the formation of an inflammatory reactive environment causing cell death which in turn is responsible for the spread of neurological deficits and the functional losses but which also leads to formation of the spinal scar and triggers regenerative mechanisms [33]. Axonal regrowth and restoration of functional network are challenged by the limited plasticity of the spinal cord due to a dense fibrotic scar and an environment rich in pro-inflammatory cytokines, myelin fragments, chondroitin sulfate proteoglycans (CSPGs), damage-associated molecular patterns (DAMPs) and other inhibitory molecules [5,34]. Conversely, axonal regrowth and synaptic reestablishment need the presence and the secretion of anti-inflammatory cytokines, growth factors and permissive molecules which are at least partly supplied by ependymal cells [35,36]. Neuronal survival and connectivity are also modulated by the perineuronal net (PNN). Indeed, at early stage (14 days after SCI), this structure, which surrounds the soma of neurons and their dendritic processes, provides protection against inflammation and apoptosis-induced molecules. Then, at later stage (56 days after SCI), it is gradually degraded, mediated by microglial cells, to allow new synaptic connections [37,38].

The inflammatory response is not limited to the spinal cord. Days following injury, cytokines and chemokines are detectable in rodent and human serum [8], and consequences associated with peripheral immune cell activation are present in other organs such as the spleen [39], kidneys, lungs [40], liver [41], adipose tissue [42] and brain [43]. For example, SCI is associated with brain inflammation and cognitive deficits, particularly in the OB, cortex, thalamus and hippocampus. Recent studies report that the spread of inflammation is due to the presence of cytokines and immune cells into cerebrospinal fluid after SCI [44].

Therefore, we asked whether SCI could impair the known effects of cell therapy: bOEC transplantation. The bOEC transplantation effects are well described in several species including dogs, cats, rodents and non-human primates after SCI as well as in humans [45]. After SCI, bOEC transplantation improves functional recovery by modulation of spinal scar composition, both from a cellular and molecular levels. Analyses of molecules secreted by bOECs show their ability to produce *in vitro* and *in vivo*, growth factors, ECM modulating molecules and axonal regrowth facilitating molecules [45]. When transplanted after SCI, bOECs participate in neuroprotection, limiting secondary injury and improving the preservation of axonal networks spared by primary damages [32]. Through metalloproteinase secretion, bOECs degrade the axonal inhibitory ECM, in particular CSPGs present in the spinal scar and stimulate the production of axonal regrowth permissive molecules [28,46,47]. They also modulate microglial/macrophages activation through direct communication, through the release of cytokines and chemokines and by indirect effects in sparing spinal cord tissue. They help to reduce microglial/macrophages activation, allowing permissive tissue for neuronal network regeneration [48]. Overall, combined effects of bOEC transplantation lead to improve spinal cord plasticity allowing axonal regrowth, myelination and stabilization of neuronal networks favoring functional recovery [49].

However, it has also been reported that cell transplantation can induce pain-related behavior such as allodynia [29]. This can be explained by the release of growth factors that stimulate neuronal survival and by the enhancement of axonal regrowth. Moreover, it can be due to an overactivity of the preserved axons, induced by the cellular environment and possibly by an increase in plasticity [50]. Indeed, to reconstitute neuronal networks, PNN surrounding neurons must be modulated over time to allow synaptic connections. It has recently been reported that reduction of PNN contributes to neuronal overactivity which in turn induces pain [51].

Here, our results provide new informations on the mechanisms induced by bOEC transplantation after SCI. Indeed, we show that bOEC transplantation promotes the formation of axon permissive spinal scar by enhancing laminin deposition (Fig. 6E-F). In addition, to its effects on spinal scar, bOEC transplantation favors axonal regrowth and axon sprouting into the lesion core (Fig. 6I and J). Furthermore, our results demonstrate also that bOEC transplantation improves synaptic connectivity, objectified by VGlut1+/BDA+ co-staining, at the chronic stage (56 days after SCI), at rostral part of the spinal cord and at the lesion core (Fig. 6L). Interestingly, results regarding synaptic formation do not correlate with a decrease of PNN density at later stage (Fig. 7H, I, K and L). Nevertheless, we can observe that the PNN is preserved and increased at early stage after transplantation compared to untreated injured mice (Fig. 7F). We can hypothesize that, preservation of PNN protects neurons and increases their survival during the acute phase and the beginning of the chronic phase (28 days after SCI) (Fig. 7D and G). Altogether, bOECs abilities contribute to the regeneration of axonal networks and the formation of new synapses, both improving functional motor recovery (Fig. 4), but are also responsible of pain related thermal stimuli (Fig. 6A). It is important to note that the identification of an increase in the number of synapses does not indicate that the network is functional. Electrophysiological analyses would allow us to highlight the functionality of synapses by examining neuronal connectivity between the neuron and its target.

Altogether, our results confirm the known effects of bOEC transplantation previously described in other studies, regarding expression of molecules modulating spinal scar and axonal regeneration. Moreover, here we described the main role played by OECs on microglia/macrophages reactivity and polarization. In fact, our results show that bOEC transplantation can modulate the ratio of pro-inflammatory (CD86+) and anti-inflammatory (CD206+) microglia/macrophages present into the lesion site. We have also deepened the knowledge about bOECs effects on modulation of the PNN, which can, at least in part, explain the increase of painful sensitivity observed in bOECs H group.

### Therapeutic effects of bOEC transplantation are impaired by SCI in an autograft model

To determine whether SCI can modulate therapeutic effects of cells in autologous transplantation paradigm, we have transplanted bOECs, obtained from SCI mice and collected 7 days after the lesion, in other SCI mice. Our findings demonstrate for the first time that the initial trauma reduces therapeutic effects of bOECs. In fact, bOEC (from injured mice) transplantation, shows no significant difference in terms of functional recovery compared to SCI (untreated animals) and bOECs H (bOECs obtained from uninjured mice) groups. On the contrary, it shows a trend with an intermediate effect and a high heterogeneity in functional performances (Fig. 4C-K), as well as in hindpaw placements during locomotion (Fig. 4M and N). This potential difference in term of neuroregenerative properties was confirmed by the assessment of spinal scar composition. Indeed, transplantation of bOECs I does not change the composition of the spinal scar compared to untreated mice. While bOEC H transplantation reduces the overall size of the fibrotic scar and thus reduces the area without astrocytes, bOEC I transplantation fails to do the same (Fig. 5B, C, F, G, J and K). Analyses of the demyelinated areas also show that bOEC H transplantation reduces MBP negative areas 14 and 56 days after SCI whereas bOEC I transplantation do not reduce it 14 and 28 days after SCI but could only decrease it 56 days after SCI (Fig. 5D, H and L respectively).

Thus, we hypothesized that the modulation of the spinal scar may be related to differences between bOECs H and bOECs I survival after SCI and transplantation. Our results show that the survival of bOECs H and I after transplantation are identical. In fact, 4 weeks after SCI, based on two different approaches, we could not find any surviving cells in the spinal cord in either group of bOECs transplanted (Fig. 3). These results are of primary importance because they show that the differences observed after bOEC H and bOEC I transplantation are not due to an effect on cell survival but are mainly due to intrinsic differences between cells *per se*.

To confirm this hypothesis, we examined the effects of bOEC I transplantation on laminin deposition, axonal regrowth and synapse formation. Our results show that, overall bOEC I transplantation decreases laminin deposition compared to bOEC H transplantation (Fig. 6E and F), has limited effects on axonal regeneration resulting in a decrease of vGlut1 synapse formation without major influences on axonal sprouting (Fig. 6I-L). In addition to these experiments, neuronal survival and PNN were assessed after bOEC I transplantation. It appears that bOEC I transplantation does not significantly improve neuronal survival after SCI compared to untreated animals whereas bOEC H transplantation improves it 14 and 28 days after SCI (Fig. 7 A, B, C, D, G and J). More interestingly, WFA analysis shows that bOEC I transplantation enhances the ratio of WFA+/NeuN+ neurons 56 days after SCI (Fig. 7K). As described above, it has been reported that PNN degradation is a necessary process after injury to induce neuronal plasticity and the establishment of synaptic connections at the chronic stage [38], which may be related to the fact that bOEC I transplantation fails to enhance formation of new synapses 56 days after SCI (Figure 6L).

These results could be explained by the differences in gene expression observed between bOEC H and bOEC I primary cultures in our qRT-PCR experiments (Fig. 2G). In fact, bOEC I primary cultures overexpresses genes involved in scar forming molecules, such as *ctgf*, *fgf1*, *tgf*β*2* or *3*, which stimulate proliferation of stromal cells and type A pericytes and also increase ECM deposition by astrocytes, leading to the generation of a larger fibrotic scar. This analysis also demonstrate that other gene expressions are modulated in bOEC I primary cultures. Indeed, a set of genes implicated in ECM modulation and axonal plasticity is upregulated as *sema3f*, *neo1*, *ntn1* or *lgals1* (Fig. 2G) which are known to negatively influence axonal regrowth.

Altogether, these results suggest that the gene expression changes observed in bOEC I primary cultures directly affect spinal scar formation, neuronal survival and axonal regrowth and reduce the therapeutic efficacy of these cells.

As described above, microglia/macrophage populations are important players in spinal cord regeneration. The number and polarization of these cells have a strong influence on spinal tissue scarring and axonal regeneration [52]. We found that *in vitro*, microglial cell cultures (BV2 cell line) are modulated by the molecules secreted by bOECs, and in particular by the CM of bOEC I primary cultures. In fact, BV2 cultured with bOECs I concentrated medium, overexpress genes encoding cytokines, chemokines, trophic factors and cellular receptors. Among these modulated genes, most of them are involved in microglial activation (e.g., *csf1*, *cx3cr1*, *p2rx4*, *p2r12*) and in polarization towards a pro-inflammatory phenotype (e.g., *ccl5*, *il1*β, *cxcr4*, *ifngr2*), while genes expression of BV2 cultured with bOECs H concentrated medium is closer to the gene expression of BV2 control cultures (without CM) (Fig. 8K). BV2 cultured with bOECs H concentrated medium down regulate the expression of pro-inflammatory genes compared to BV2 control while at the same time BV2 bOECs I upregulate their expression of pro-inflammatory genes compared to control (Fig. 8L and M). These results can be linked to the increase in *il1*β and *tnf-*α gene expression observed in bOECs I primary culture in our qRT-PCR experiments (Fig. 2G). At the opposite, *in vivo* analyses, show that there is no major difference in microglia/macrophage activation and polarization between bOECs H and bOEC I transplantation at the early stage and at the beginning of the chronic stage after SCI (Fig. 8A-G). Nevertheless, it can be noted that only the bOECs H group shows a significant increase in the percentage of pro-regenerative/anti-inflammatory microglia/macrophages, identified by CD206, in the rostral and caudal parts of the spinal cord compared to SCI mice at these stages. On the other hand, at the chronic stage, the time of the second microglia/macrophage wave, the percentage of pro-inflammatory microglia/macrophages is reduced after bOEC H transplantation compared to bOEC I transplantation (Fig. 8H). These results may indicate that bOECs H in the chronic stage, create a less reactive extracellular environment that favors axonal network regeneration by facilitating mechanisms involved in PNN degradation and synaptic connection formation (Fig. 6L and 7K).

Altogether, our results demonstrate for the first time the major impact of the initial injury on the therapeutic effects of cells that can be used for cellular transplantation after SCI. It also highlights one of the reasons that may explain the differences observed in spinal cord regeneration and functional recovery after cellular transplantation in mice and humans. Indeed, the initial physical trauma does not only lead to localized disturbances at the site of the injury [41]. The whole body is in an “inflammatory state” in response to circulating cytokines, chemokines and immune cells, causing intrinsic changes in the cells used for autografting that are responsible for their reduced therapeutic efficacy. The inflammatory response that occurs after SCI is the main actor in the propagation of trauma. The use of anti-inflammatory drugs to restrict inflammation in the first hours, days and weeks after SCI, in order to reduce the cellular response and increase the therapeutic effect of autologous cells, may have adverse effects. Indeed, it has been shown that the immune response, and in particular the microglial/macrophage response, is important for tissue regeneration after SCI. The use of anti-inflammatory drugs has been shown to worsen spinal scar size, reduce axonal regeneration and limit functional recovery [53,54]. In our study, we chose to collect the OBs 7 days after SCI, at early stage. One solution would be to collect cells several weeks or months after SCI when immune response is lower, but it is important to consider the chronology of the physiopathology. It has been described, at least in animal studies, that the most effective therapeutic window is at early stage after SCI. Indeed, the use of an effective therapy at early stage should make it possible to reduce axonal and, more broadly, tissue damages and limit the exacerbation of the inflammatory response.

## Conclusion

Finally, our results should be put into a clinical perspective. Indeed, autograft transplantation is considered as potential treatment to cure SCI. However, our results demonstrate for the first time that traumatic SCI impairs the neuroregenerative properties of cells used as therapy when obtained from injured animals. It is reasonable to assume that this will also be the case in human patients suffering from spinal cord injuries, which would call into question the feasibility and therapeutic efficacy of these transplants.

